# Sex and time-of-day differences in kidney oxygen consumption

**DOI:** 10.1101/2023.04.18.537340

**Authors:** Pritha Dutta, Anita T. Layton

## Abstract

Renal hemodynamics, renal transporter expression levels, and urine excretion all exhibit circadian variations. Disturbance of these diurnal patterns is associated with hypertension and chronic kidney disease. Renal hemodynamics determines oxygen delivery, whereas renal transport and metabolism determines oxygen consumption. The balance between oxygen delivery and consumption then yields renal oxygenation. We hypothesized that kidney oxygenation also demonstrates 24-h periodicity. Another notable modulator of kidney function is sex, which has impacts on renal hemodynamics and transport function that are regulated by as well as independent of the circadian clock. The goal of this study was to investigate the diurnal and sexual variations in renal oxygen consumption and oxygenation. For this purpose, we developed computational models of rat kidney function that represent sexual dimorphism and circadian variation in renal hemodynamics and transporter activities. Model simulations predicted substantial differences in tubular Na^+^ transport and oxygen consumption among different nephron segments. We also simulated the effect of loop diuretics, which are used in the treatment of renal hypoxia, on the outer medullary oxygen tension. Our model predicted a significantly higher effect of loop diuretics on renal oxygenation in female rats compared to male rats. In addition, loop diuretics were more effective when administered during the active phase.

## 1. Introduction

The kidneys are responsible for maintaining chemical and fluid homeostasis in the body by removing waste products and excess solutes and fluids. Hence, the kidneys receive a high blood flow, approximately 25% of the cardiac output. To process that amount of blood, the kidneys consume the second highest amount of oxygen, normalized by organ weight, after the heart. However, compared to other organs, renal oxygen extraction is very low, approximately 10-15%, whereas that for the heart is about 45%. Thus, kidneys are particularly susceptible to hypoxia, which plays an important role in the development of acute or chronic kidney diseases [1-5].

The kidneys reabsorb nearly 99% of the filtered Na^+^. The proximal tubule reabsorbs approximately 60-70% of the filtered Na^+^ followed by the thick ascending loop of Henle which reabsorbs about 25-30%; the distal tubules reabsorb only 10% of the filtered Na^+^ [6]. Na^+^ reabsorption is the primary energy consuming process in the kidney driven by the sodium-potassium pump Na-K-ATPase on the basolateral membrane. Na-K-ATPase transports 3 Na^+^ out of the cell in exchange for 2 K^+^ with the cost of hydrolyzing 1 ATP molecule [7]. To retrieve nearly 99% of filtered Na^+^, a continuous supply of large amount of ATP is required. Hence, sufficient oxygen delivery to the kidneys is required to meet this ATP demand.

Renal blood flow in the kidneys is highly heterogeneous; the cortex is well perfused but the medulla receives only 10-15% of the renal perfusion. The medullary perfusion is low, in part, to maintain osmotic gradients and urine concentrating ability of the medulla [8]. However, the medullary oxygen consumption accounts for approximately 20% of the renal oxygen consumption. The medullary thick ascending limbs have high oxygen requirement for Na^+^ transport but low oxygen delivery which makes this segment particularly prone to hypoxia [8-10]. The oxygen shunting between the descending and ascending vasa recta in the medulla also contributes to the low oxygen availability in this region [8, 10]. Additionally, tubular cells within the S3 segment of the proximal tubule, which is in the outer medullary region, have high oxygen demand due to the abundance of active Na^+^-K^+^-ATPases. The combination of these factors can aggravate outer medullary hypoxia even in healthy kidneys [2, 8, 10, 11].

There is a close relation between glomerular filtration rate (GFR), renal Na^+^ reabsorption, and oxygen consumption. In most organs, an increased demand of oxygen is met by an increased blood flow to increase oxygen delivery. However, an increased renal blood flow simultaneously increases tubular Na^+^ load due to increased GFR. Since sodium reabsorption is the major renal oxygen consuming process, increased renal blood flow also increases the oxygen demand. Renal oxygen extraction remains stable over a wide range of renal blood flow, indicating counteraction of increased oxygen delivery by increased oxygen consumption [12].

In recent years, two new dimensions have emerged for investigation of kidney function: sex and time of day. Like nearly every tissue and organ, the structure and function of the mammalian kidney is regulated by sex hormones [13, 14]. In addition to organ size, the abundance patterns of membrane transporters and channels in rodent kidneys have been reported to be sexually dimorphic [15]. For instance, female rat proximal tubules exhibit lower Na^+^/H^+^ exchanger 3 (NHE3) abundance and activity compared to the male counterparts and thus reabsorb a substantially lower fraction of the filtered Na^+^. The higher fractional Na^+^ distal delivery in females is handled by the augmented transport capacity in the downstream distal tubular segments, particularly by the Na^+^-K^+^-Cl^−^ cotransporter 2 (NKCC2) and Na^+^-Cl^−^ cotransporter (NCC) transporters in the thick ascending limbs and distal convoluted tubules. Female rats exhibit higher NKCC2 and NCC activity (almost double) relative to males. Male rats transport a larger fraction of the filtered Na^+^ through NHE3 in the proximal tubules, whereas female rats transport a larger fraction of the filtered Na^+^ through NKCC2 in the medullary thick ascending limbs [15-18]. Thus, cortical and medullary oxygen consumptions are different between the sexes. In addition, different segments have different transport efficiency-a segment having higher paracellular transport would transport more Na^+^ moles per mole of O_2_ consumed [19, 20]. Hence, a goal of this study is to investigate the sex differences in whole-kidney and regional oxygen consumption and segmental transport efficiency, which we hypothesize would arise from the difference in segmental distribution of Na^+^ transport between males and females.

Kidney function is regulated by the circadian clock, such that GFR, filtered electrolyte loads, urine volume, and urinary excretion exhibit significant diurnal rhythms [21-23]. These diurnal rhythms occur partly due to the regulation of renal transporter proteins, including NHE3, NKCC2, NCC, and epithelial Na^+^ channels (ENaC) by clock proteins brain and muscle ARNT-like 1 (BMAL1) and period circadian regulator 1 (PER1) [24-27]. Interestingly, sex and time-of-day are not two independent regulators of kidney function. For instance, notable sex differences have been reported in the regulation of renal Na^+^ transport by BMAL1 [28]. Thus, any investigation of kidney metabolism and function must take into account sex and time of day as variables.

To assess the sex- and time-of-day specific differences in oxygen consumption along different nephron segments, we developed computational models of circadian regulation of solute and water epithelial transport in male and female rats. We have previously developed epithelial transport models of the rat kidney that are sex specific [16-18, 29, 30] or that incorporate circadian rhythms in GFR and transport activities [31, 32], but none that considers both variables. Using the sex- and time-of-day specific models, we simulated the circadian and sexual variation in Na^+^ transport and oxygen variation along different nephron segments and regions (cortical and medullary regions). We also developed an equation to compute renal oxygen tension based on the model’s predicted oxygen consumption. Using this equation, we predicted the change in outer medullary oxygenation on treatment with loop diuretics, which are commonly used to treat hypoxia, to evaluate the difference in effectiveness in male and female rats.

## 2. Methodology

We developed sex- and time-of-day specific models of the rat kidney function. A schematic diagram of the various cell types is given in Fig. 1. The models are based on the epithelial cell-based model of solute transport in a rat kidney developed by Layton et al. [16, 33]. These models represent the sexual dimorphism in tubular dimensions, single nephron glomerular filtration rate (SNGFR), and expression levels of apical and basolateral transporters in rats. The models also represent the circadian regulation of GFR and transporter activities in a light-dark cycle. GFR and selected transporter activities are set to vary as sinusoidal functions of time. Model parameters that exhibit circadian rhythms are summarized in Table 1 and Fig. 2.

**Table 1.**
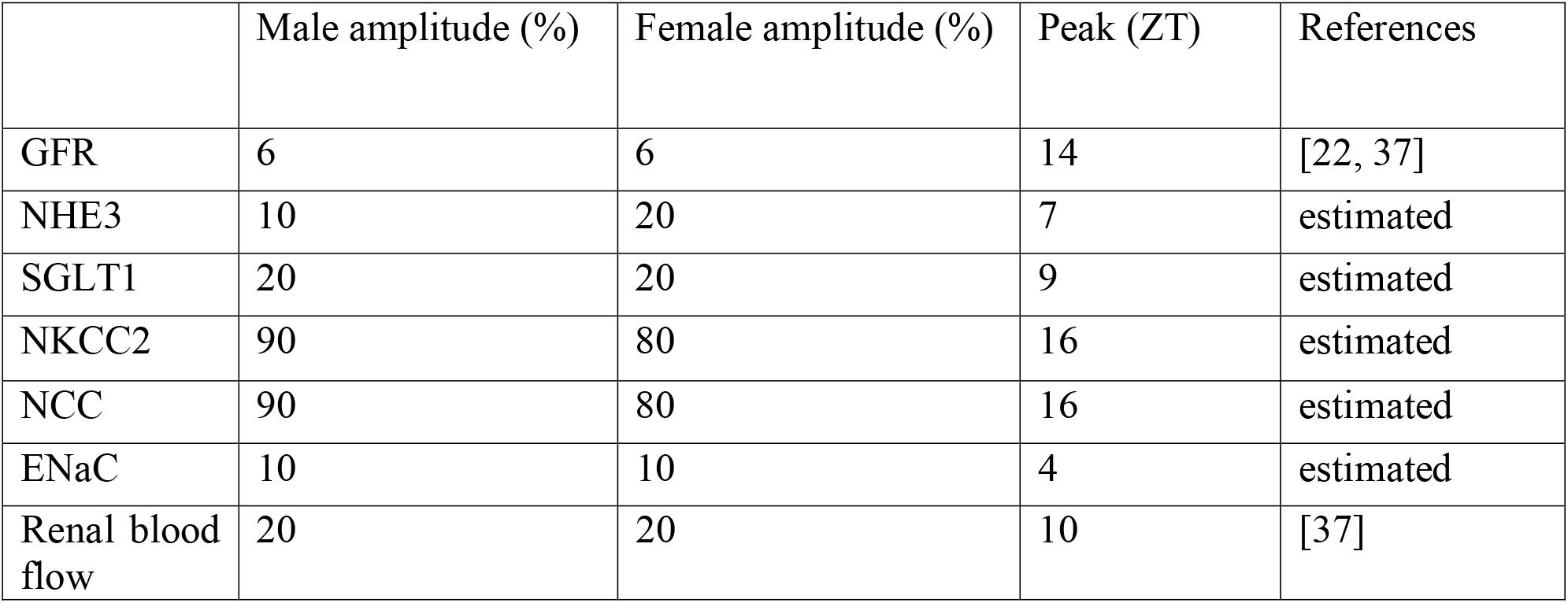
Peak times and oscillation amplitudes of model parameters that exhibit circadian rhythms.

|  | Male amplitude (%) | Female amplitude (%) | Peak (ZT) | References |
| --- | --- | --- | --- | --- |
| GFR | 6 | 6 | 14 | [22, 37] |
| NHE3 | 10 | 20 | 7 | estimated |
| SGLT1 | 20 | 20 | 9 | estimated |
| NKCC2 | 90 | 80 | 16 | estimated |
| NCC | 90 | 80 | 16 | estimated |
| ENaC | 10 | 10 | 4 | estimated |
| Renal blood flow | 20 | 20 | 10 | [37] |

**Figure 1.**
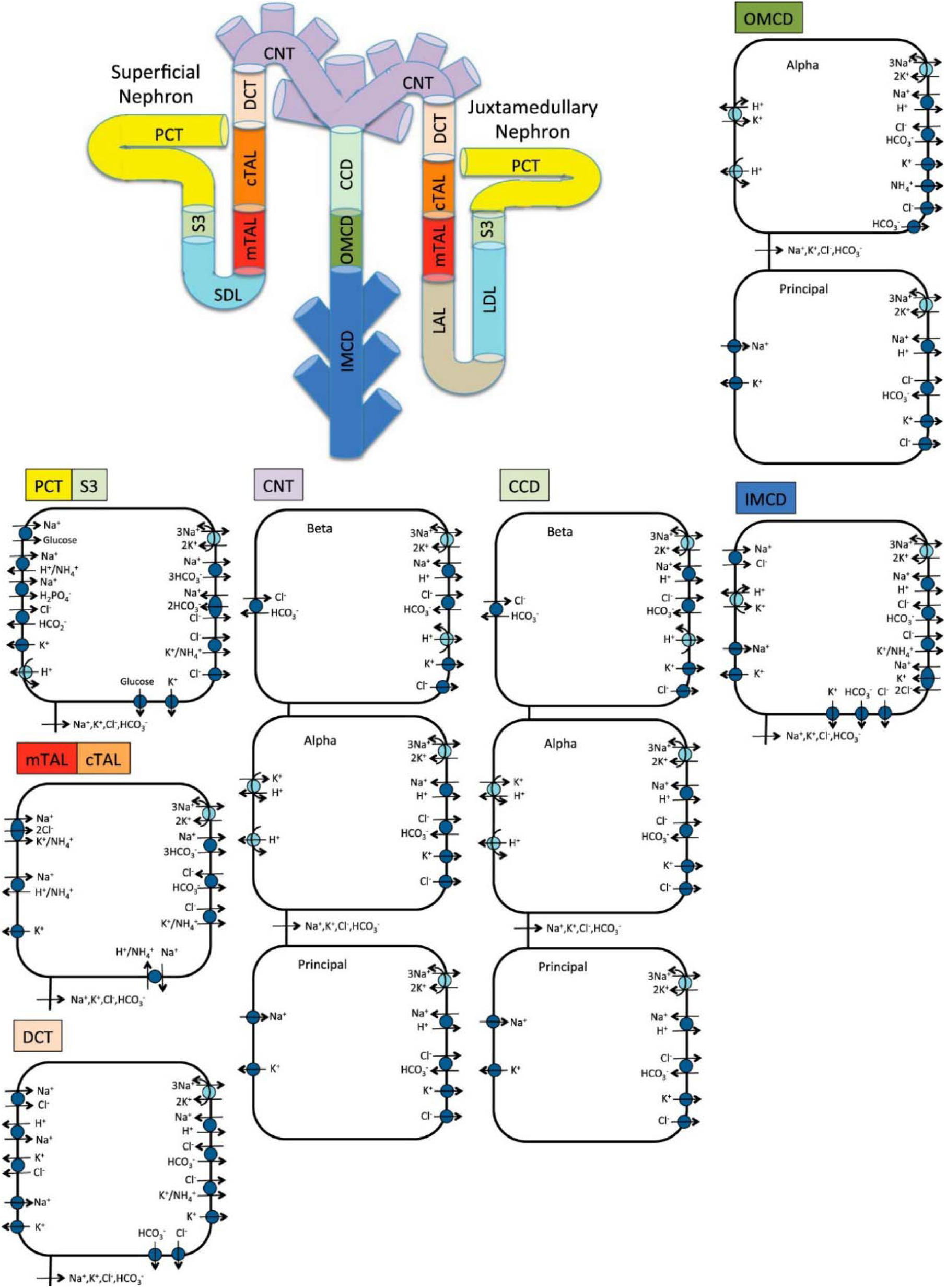
Schematic diagram of the nephron system (not to scale). The model includes one representative superficial nephron and five representative juxtamedullary nephrons, each scaled by the appropriate population ratio. Only the superficial nephron and one juxtamedullary nephron are shown. Along each nephron, the model accounts for the transport of water and 15 solutes. The diagram displays only the main Na^+^, K^+^, and Cl^−^ transporters. PCT, proximal convoluted tubule; SDL, short descending limb; mTAL, medullary thick ascending limb; cTAL, cortical thick ascending limb; DCT, distal convoluted tubule; CNT, connecting tubule; CCD, cortical collecting duct; OMCD, outer-medullary collecting duct; IMCD, inner-medullary collecting duct; LDL/LAL, thin descending/ascending limb. Adopted from [33].

**Figure 2.**
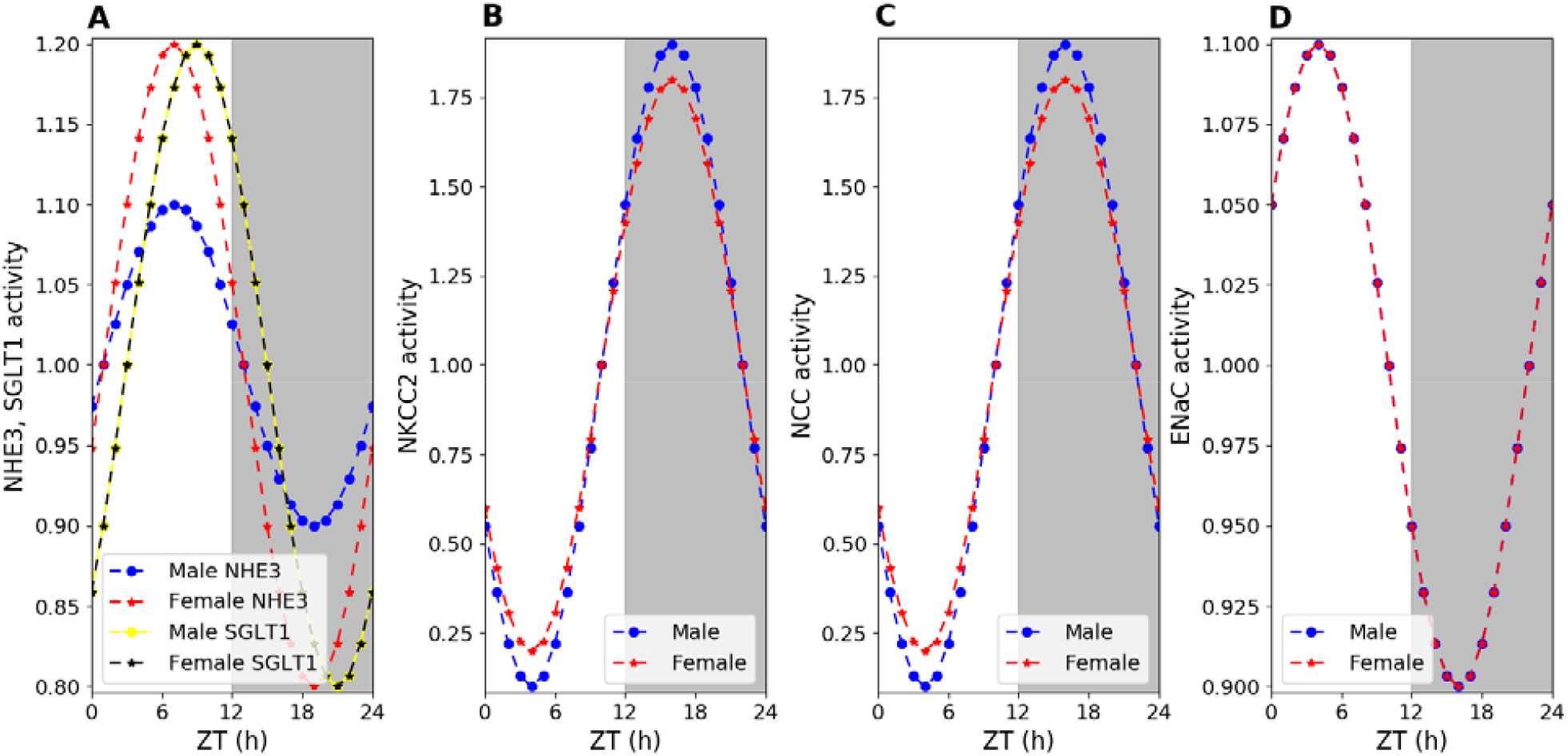
Time profiles of circadian regulated transporter activities. (A) Na^+^/H^+^ exchanger 3 (NHE3) and sodium-glucose transporter 1 (SGLT1) activities; (B) Na^+^-K^+^-Cl^−^ cotransporter 2 (NKCC2) activity; (C) Na^+^-Cl^−^ cotransporter (NCC) activity; (D) epithelial sodium channel (ENaC) activity. Values are normalized to mean values.

### 2.1 Model structure

The model for male or female rat kidney includes six classes of nephrons: a superficial nephron and five juxtamedullary nephrons that are assumed to reach depths of 1, 2, 3, 4, and 5 mm into the inner medulla. The superficial nephron is assumed to account for 2/3 of the nephron population, while the juxtamedullary nephrons account for 0.4/3, 0.3/3, 0.15/3, 0.1/3, and 0.05/3 of the nephron population. The superficial nephron includes the proximal tubule, short descending limb, thick ascending limb, distal convoluted tubule, and connecting tubule segments. The juxtamedullary nephrons include all the segments of the superficial nephron with the addition of the long descending limbs and ascending thin limbs; these are the segments of the loops of Henle that extend into the inner medulla. The length of the long descending limbs and ascending limbs are determined by the type of juxtamedullary nephron being modeled. The connecting tubules of the five juxtamedullary nephrons and the superficial nephron coalesce into the cortical collecting duct. The SNGFR is set to ∼30 and ∼45 nl/min for the superficial and juxtamedullary nephrons, respectively, in male rats and to ∼24 and ∼36 nl/min for the superficial and juxtamedullary nephrons, respectively, in female rats.

Each nephron segment is modeled as a tubule lined by a layer of epithelial cells in which the apical and basolateral transporters vary depending on the segment and cell type (intercalated and principal cells). A paracellular pathway exists between neighboring cells. The models account for the following 15 solutes: Na^+^, K^+^, Cl^-^, HCO_3_^-^, H_2_CO_3_, CO_2_, NH_3_, NH_4_^+^, HPO_4_^2-^, H_2_PO_4_^-^, H^+^, HCO_2_^-^, H_2_CO_2_, urea, and glucose. Male and female models differ in parameters describing membrane transporter and channel activities and paracellular permeabilities. The models consist of a large system of coupled ordinary differential and algebraic equations, which impose mass conservation and electroneutrality, and calculate transmembrane and paracellular fluxes [34]. Water fluxes are driven by osmotic and hydrostatic pressure differences. Transmembrane solute fluxes may include passive and active components. An uncharged solute may be transported across a membrane passively by a concentration gradient, whereas a charged solute may be transported by an electrochemical potential gradient across an ion channel. Additional components may include coupled transport across co-transporters and/or exchangers and primary active transport across ATP-driven pumps, the activities of which may exhibit circadian rhythms [24-27]. The model predicts the steady-state values of urine volume, urinary excretion rates of individual solutes, as well as luminal fluid flow throughout the nephron, hydrostatic pressure, membrane potential, luminal and cytosolic solute concentrations, and transcellular and paracellular fluxes through transporters and channels [16].

### 2.2 Circadian rhythms in transport parameters

We estimated the male rat transporter activity amplitudes from the diurnal variations in water, sodium, and potassium excretion rates observed in male Sprague Dawley rats [35]. Though female rats have lower water, sodium, and potassium excretion rates relative to male rats, the rates are not significantly different [15]. Hence, we assumed that the excretion rates of female rats are 10% lower than those of male rats. We used the root mean squared error between the simulated and experimental excretion rates to fit the transporter activity amplitudes. The transporter activity peaks were adapted from [36] based on the GFR values. In [36], GFR peaks at ZT16, whereas according to the rat data [22, 37] GFR peaks at ZT14. Based on this, we shifted the transporter activity peaks. The experimental and simulated volume, Na^+^, and K^+^ excretion rates for male and female rats are given in Fig. S1.

In a light-dark cycle, ZT0 (lights on) marks the start of the rest phase for nocturnal animals, while ZT12 (lights off) denotes the start of the active phase. The model represents the circadian rhythm in transporter activities (e.g., NHE3, NKCC2, etc., see Table 1) as

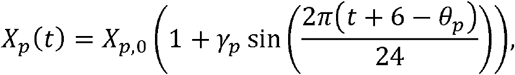

where *t* is the Zeitgeber time (ZT), *X*_*p,o*_ represents the average activity, *γ*_*p*_ denotes the fractional oscillation amplitude, and *θ*_*p*_ denotes the peak time. The circadian rhythms in GFR are represented similarly. Parameter values are specified in Table 1.

To determine model prediction at a given ZT (e.g., 14), GFR and transporter activities were set at their ZT = 14 values. The steady-state model solution was then computed. The use of a steady-state solution for a model with circadian rhythm is justified by the much shorter timescale of tubular flows compared with circadian oscillations.

### 2.3. Oxygen consumption along the nephron

Oxygen consumption (Q_02_) consists of two parts: 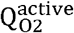, which represents the oxygen consumed to actively reabsorb Na^+^, and 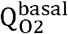,, which represents the oxygen consumed for other transport processes and intracellular biochemical reactions [33, 38]. 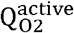, is calculated based on the ATP consumption of basolateral Na-K-ATPase pumps. Na-K-ATPase pumps require 1 mole of ATP to pump out 3 moles of Na^+^, and oxidative metabolism yields about 5 moles of ATP per mole of O_2_ consumed [39]. Thus, 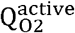, is determined as

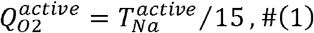

where 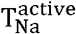, is the rate of Na^+^ transport across Na-K-ATPase pumps [33, 38].

In rats, the whole kidney basal-to-total Q_02_ ratio has been estimated as 25–30% [40]. We assumed that 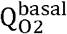 is fixed and equal to 25% of the total Q_02_ under baseline conditions [33, 38], such that,

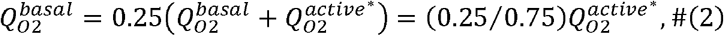

where * denotes baseline conditions.

The efficiency of oxygen utilization can be evaluated by computing the number of moles of Na^+^ reabsorbed per mole of O_2_ consumed [33, 38]:

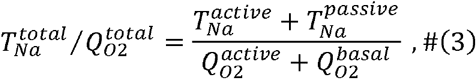

where 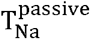, denotes the rate of passive Na^+^ reabsorption.

### 2.4 Estimation of partial pressure of oxygen (p_02_)

The renal outer medullary partial pressure of oxygen (*p*_*02*_, mmHg) is estimated using the following equation:

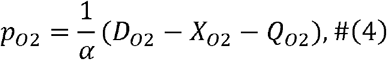

where *D*_*0*2_ denotes oxygen delivery (µmol/min) from the descending vasa recta, *X*_*0*2_ denotes oxygen shunting (µmol/min) between the descending and ascending vasa recta, *Q*_*0*2_ denotes the oxygen consumption in the renal outer medulla (µmol/min), and α represents the pressure to µmol conversion factor (µmol/min/mmHg).

Oxygen delivery, *D*_*0*2_, is calculated as [41]

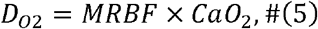

where *MRBF* denotes medullary renal blood flow (mL/min) and *CaO*_*2*_ denotes the arterial oxygen content. We assumed the mean *MRBF* for male rats to be 2.26 mL/min (assuming a mean kidney weight of 1.189 g) [42, 43]. In the model, the single nephron GFR for male rats is assumed to be 25% higher than that for female rats. Hence, we assumed the mean *MRBF* for female rats to be 1.81 mL/min. The arterial oxygen content is calculated as [41]

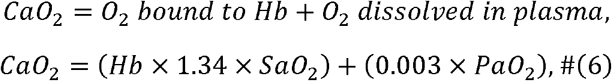

where *Hb* denotes the amount of hemoglobin which is 146 and 141 g/L in male and female rats, respectively [44]. Amount of oxygen carried by 1 g of Hb is 1.34 mL [41]. *SaO*_*2*_ denotes the arterial oxygen saturation which we assume as 95%. Oxygen dissolved in plasma is 0.003 mL O_2_/ 100 mL blood/ mmHg [41]. *PaO*_*2*_ denotes the arterial partial pressure of oxygen which is ∼88 mmHg in rats [45].

Approximately 2.6% of the total oxygen delivered through the descending vasa recta is shunted to the ascending vasa recta [46]. Hence,

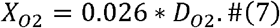

Mean *p*_O2_ in outer medulla is 15-30 mmHg [8]; we assumed this as 22.5 mmHg. We used the peak time and oscillation amplitude of the outer medullary *p*_O2_ as 13 h (ZT) and 8%, respectively [47]. We assumed the same medullary *p*_O2_ for male and female rats [30].

### 2.4 Simulating loop diuretics

Inhibition of active transport along the loop of Henle with loop diuretics, such as furosemide, can lower oxygen consumption and increase medullary oxygen tension [48]. Loop diuretics inhibit the Na^+^ transporter, NKCC2, which is expressed on the apical membrane of the thick ascending limbs of the loops of Henle. We assumed that the loop diuretic was administrated for long enough to significantly impair the kidney’s ability to generate an axial osmolality gradient [49, 50]. Thus, we lowered the interstitial fluid concentrations of selected solutes [49]. Given that targeted deletion of NKCC2 significantly attenuates the tubuloglomerular feedback response [51], we assumed that SNGFR remained at baseline values, consistent with an experimental study in the rat [52]. Furosemide, a loop diuretic, has limited glomerular filtration and is secreted into the proximal tubular lumen by the organic anion transporter-1 (OAT1) [53]. The mRNA levels of OATs have been observed to exhibit circadian rhythms, reaching its peak in the late light phase and early dark phase [54]. This diurnal expression of OATs in the kidney may contribute to time-dependent excretion of OAT substrates such as furosemide, whose renal excretion was reduced on deletion of Bmal1 (a clock gene) in mice [54]. We represent the effect of the diurnal expression of OATs by assuming that more NKCC2 is inhibited in the dark phase (80%) compared to the light phase (70%).

## 3 Results

We conducted model simulations to predict solute and volume transport along nephron segments in male and female rats. GFR and filtered Na^+^ and K^+^ loads, which were used as inputs to predict fluid and solute flows along different nephron segments, are shown in Fig. S2. The predicted Na^+^, K^+^, Cl^−^, and volume deliveries to different nephron segments (proximl tubules, thick ascending limbs, distal convoluted tubules, connecting tubules, and collecting ducts) at different zeitgeber times in male and female rats are shown in Fig. 3. The solute and volume deliveries show a diurnal rhythm, peaking in the dark (active) phase, and is in phase with the GFR. The rat model predicted that more than half of the filtered Na^+^, K^+^, Cl^−^, and volume are reabsorbed along the proximal tubules with the thick ascending limbs reabsorbing most of the remaining solutes and volume. K^+^ deliveries to the connecting tubules and collecting ducts increase because the transepithelial elctrochemical gradient causd by Na^+^ reabsorption through ENaC in the distal convoluted tubules and connecting tubules favours K^+^ secretion.

**Figure 3.**
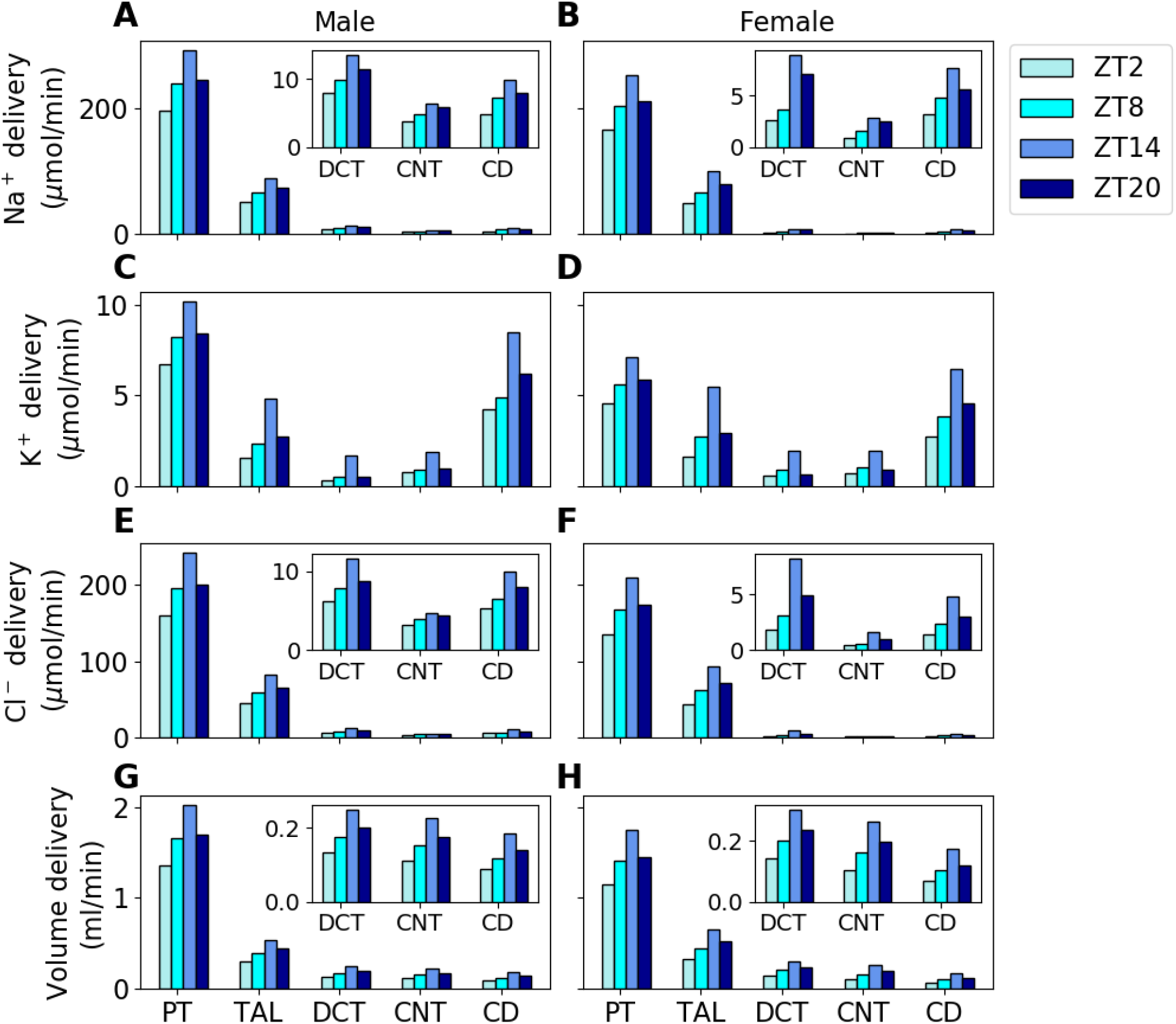
Predicted Na^+^ (A, B), K^+^ (C, D), Cl^−^ (E, F), and volume (G, H) deliveries to the proximal tubules (PT), thick ascending limbs (TAL), distal convoluted tubules (DCT), connecting tubules (CNT), and collecting ducts (CD) of male and female rats at zeitgeber times 2, 8, 14, and 20 h. The values are given per kidney.

### 3.1 Na^+^ transport exhibits significant sex-, time-of-day, and regional variations

The predicted (active, passive, and total) in the proximal tubules, thick ascending limbs, and distal tubules (comprising distal convoluted tubules, connecting tubules, and collecting ducts) of male and female rats at different zeitgeber times (2, 8, 14, and 20 h) are shown in Fig. 4. The predicted active, passive, and total display diurnal rhythms in phase with GFR, peaking during the active phase (ZT14). Both transcellular and paracellular T_Na_ change proportionally with luminal flow. Total T_Na_ increases by 31%, 58%, and 22% during the active phase relative to the inactive phase in the male proximal tubules, thick ascending limbs, and distal tubules, respectively. The corresponding total T_Na_ increments in the female nephron segments are 21%, 63%, and 46%, respectively. Proximal tubule active T_Na_ is higher in males compared to females because males have higher NHE3 activity and filtered Na^+^ in the proximal tubules. In addition, proximal tubule passive T_Na_ is higher in males than in females. This combined higher active and passive T_Na_ results in lower Na^+^ delivery to the male thick ascending limbs relative to females. Thus, female thick ascending limbs have higher active T_Na_ due to higher Na^+^ delivery as well as higher NKCC2 and Na^+^-K^+^-ATPase activities.

**Figure 4.**
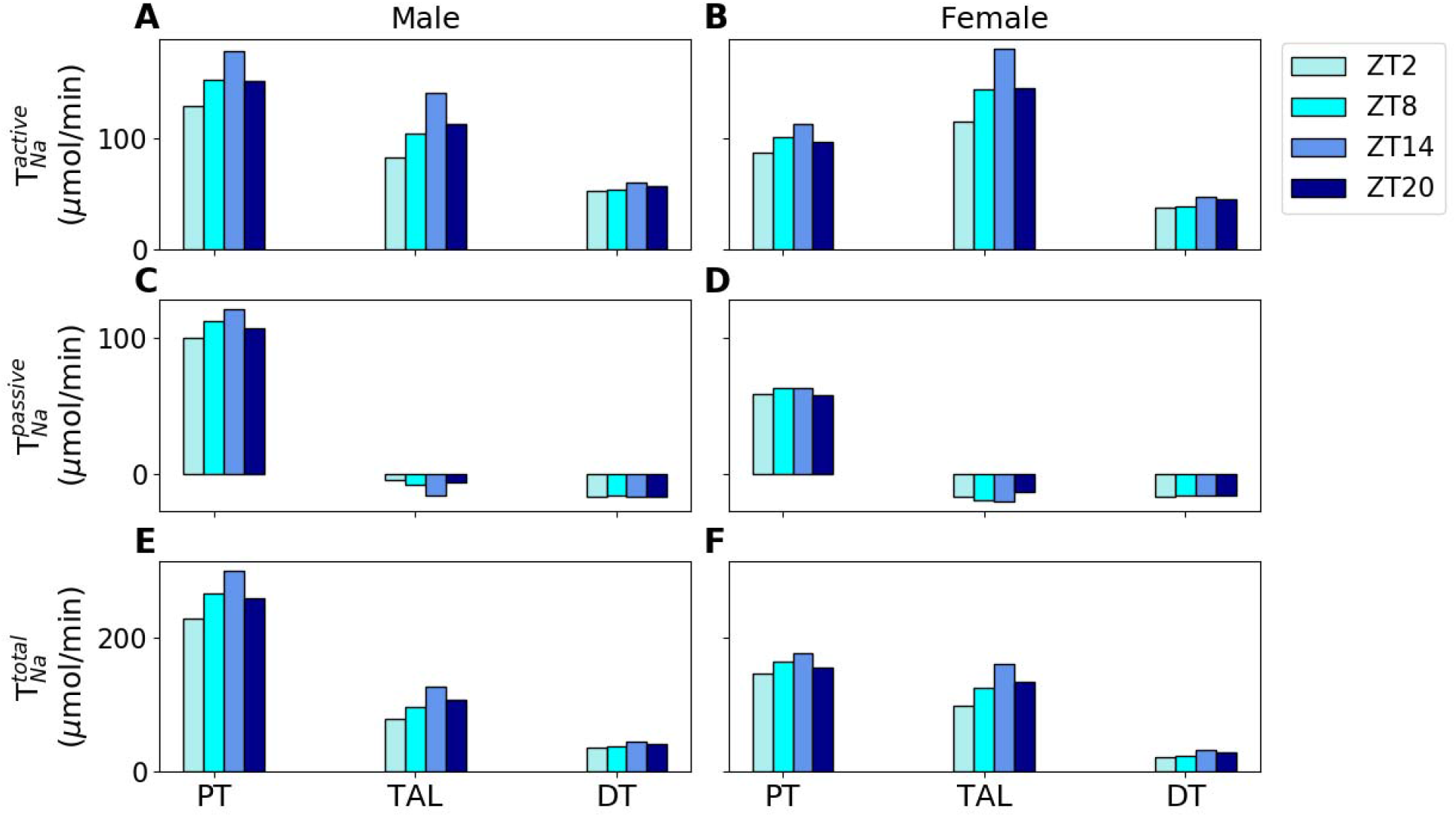
Predicted renal active (A, B), passive (C, D), and total (E, F) in the proximal tubules (PT), thick ascending limbs (TAL), and distal tubules (DT) of male and female rats at zeitgeber times 2, 8, 14 and 20 h. The values are given per kidney.

Given that the renal medulla is poorly perfused compared to the cortex, we analyzed T_Na_ for the two regions separately. Figure 5 shows the predicted T_Na_ for the cortical segments, medullary segments, and whole kidney. The predicted total T_Na_ in the cortical segments (comprising the proximal convoluted tubules, cortical thick ascending limbs, distal convoluted tubules, connecting tubules, and cortical collecting ducts) is ∼52% higher in the male rats compared to female rats. This is because in the cortical region, the majority of Na^+^ transport occurs in the proximal convoluted tubule, and male rats have higher Na^+^ transport in the proximal tubules. The predicted total T_Na_ in the medullary segments (comprising proximal straight tubule, medullary thick ascending limbs, and outer and inner medullary collecting ducts) is ∼16% higher in female rats compared to male rats. This is because in the medullary region, the majority of Na^+^ transport occurs in the medullary thick ascending limbs, and female rats have higher Na^+^ transport in the thick ascending limbs.

**Figure 5.**
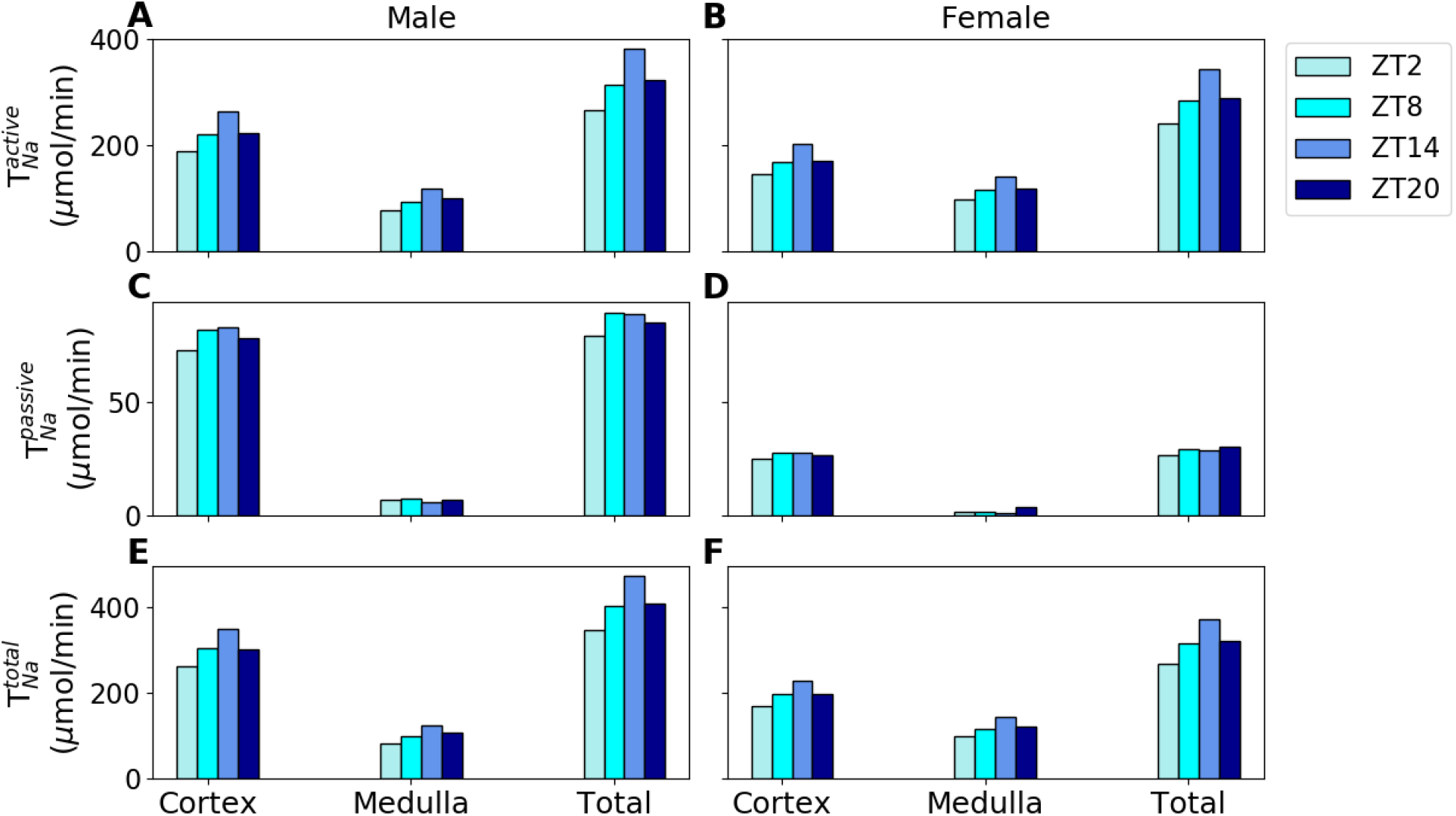
Predicted renal active (A, B), passive (C, D), and total (E, F) in the cortical segments, medullary segments, and whole kidney of male and female rats at zeitgeber times 2, 8, 14 and 20 h. The values are given per kidney.

Paracellular transport, which is passive diffusion driven by transepithelial electrochemical gradient, is the mechanism for efficient oxygen utilization as it does not require energy from ATP hydrolysis. Paracellular Na^+^ transport (passive T_Na_) in the proximal tubule follows a diurnal pattern in phase with the GFR and it is higher in the male rats due to higher Na^+^ delivery. Paracellular transport is almost zero or results in Na^+^ secretion at the thick ascending limbs due to the following reason. In the initial part of the medullary thick ascending limb, both luminal and interstitial Na^+^ concentrations are high and the positive voltage in the lumen drives Na^+^ out of the lumen into the interstitium. However, as Na^+^ is reabsorbed, the concentration gradient for Na^+^ increases which causes Na^+^ to enter the lumen from the interstitium [55].

### 3.2 exhibit significant sex-, time-of-day, and regional variations

The predicted total and efficiency of oxygen utilization () in the proximal tubules, thick ascending limbs, and distal tubules of male and female rats at different zeitgeber times (2, 8, 14, and 20 h) are shown in Fig. 6(A, B). Figure 6(C, D) shows these predicted values for the cortical segments, medullary segments, and whole kidney. Since, our model assumes passive to be constant, total changes proportionally with active. Male rats have higher oxygen consumption in the proximal tubules because of higher luminal flow and NHE3 activity, whereas female rats have higher oxygen consumption in the thick ascending limbs due to higher NKCC2 and Na^+^-K^+^-ATPase activity. Similarly, oxygen consumption in the renal cortex is ∼1/3 higher in male rats and ∼1/5 higher in female rats during both the active and inactive phases. With respect to the whole kidney, male rats have ∼10% higher oxygen consumption than female rats. Since male rat kidneys filter ∼25% more Na^+^ than the female counterparts, their whole kidney is expected to be ∼25% higher than that of female rats. However, male rats transport more Na^+^ along the proximal tubule, whereas female rats transport more Na^+^ along the thick ascending limbs. Proximal tubules have higher Na^+^ transport efficiency compared to the other nephron segments (Fig. S3) due to their high paracellular Na^+^ transport. This could be a possible reason for the lower whole kidney difference between male and female rats.

**Figure 6.**
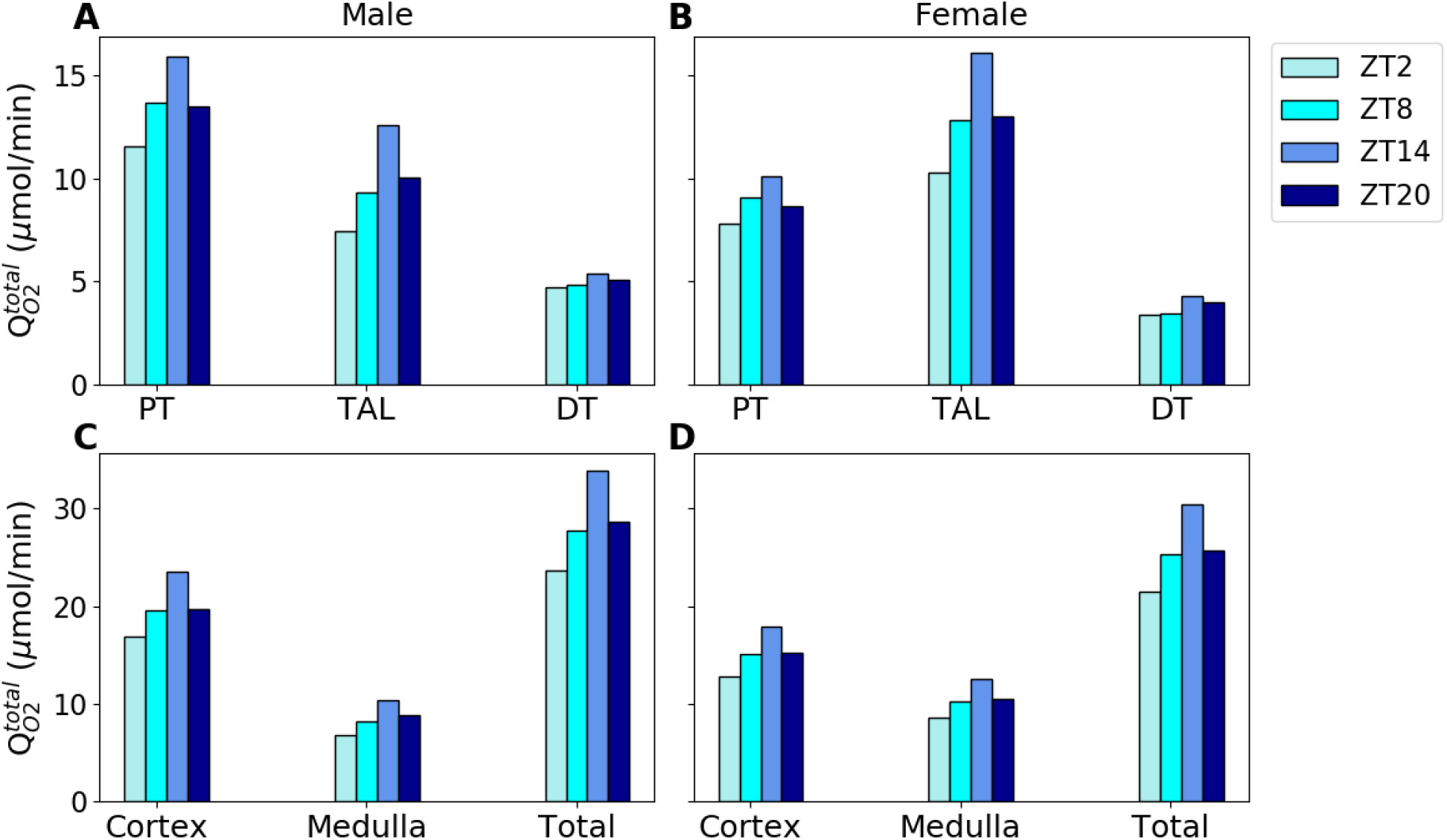
Predicted renal total in (A, B) the proximal tubules (PT), thick ascending limbs (TAL), and distal tubules (DT), and (C, D) the cortical segments, medullary segments, and whole kidney of male and female rats at zeitgeber times 2, 8, 14 and 20 h. The values are given per kidney.

The efficiency of oxygen utilization for Na^+^ reabsorption varies between tubular segments. It is higher in the proximal tubules than in the thick ascending limbs and distal tubules (Fig. S3). This is because the net paracellular transport, which does not require ATP hydrolysis and hence is an important determinant of oxygen utilization efficiency, is almost zero in the thick ascending limbs and distal tubules. Transcellular changes proportionally with luminal flow. Thus, as GFR increases, (and thus) increases at the same rate as. Thus, the model predicts that the oxygen utilization efficiency does not change significantly during the day (Fig. S3).

### 3.3 Loop diuretics have significantly greater effect on medullary oxygenation in female rats

Based on the predicted, we computed using Eq. 4. The predicted and experimental (discussed in Methodology) values are given in Fig. 7. The model predicted an ∼43% increase in during the active phase (ZT14) relative to the inactive phase (ZT2) for both male and female rats; the corresponding increase in the experimental data is ∼16%. This discrepancy is possibly due to our calculation (Eq. 5). Due to the unavailability of data on diurnal variation in, we estimated it based on the diurnal variation in renal blood flow. Renal blood flow is ∼40% higher during the active phase compared to the inactive phase [37]. This contributes to the high diurnal variation in our predicted.

**Figure 7.**
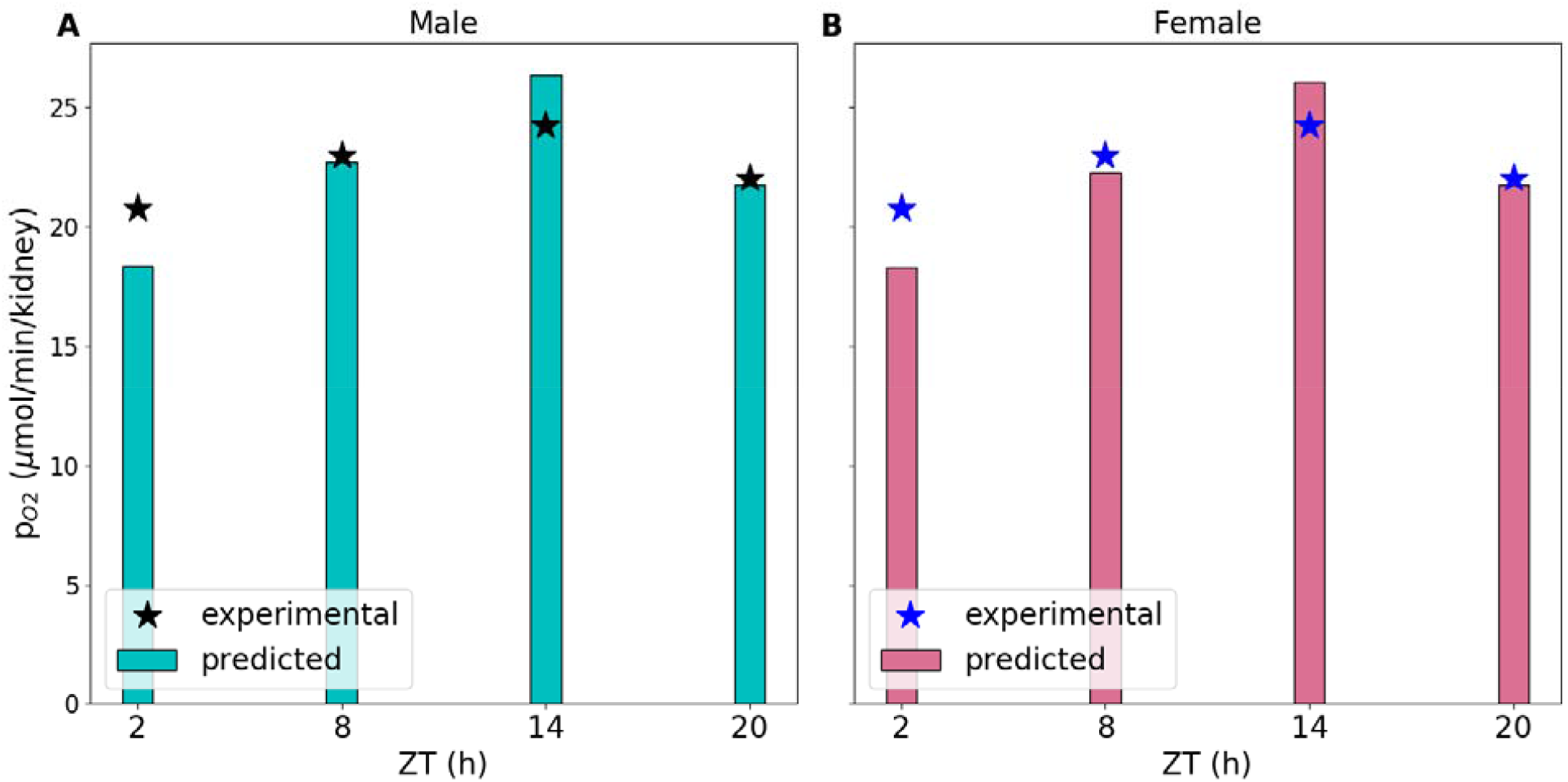
Experimental and predicted in the outer medulla of male and female rats at zeitgeber time 2, 8, 14, and 20 h.

We then used the rat models to simulate the effect of loop diuretics on oxygen consumption and renal oxygen tension. By inhibiting thick ascending limb active Na^+^ transport, loop diuretics have been found to ameliorate medullary hypoxia [48]. Figure 8 shows the regional oxygen consumption before and after intervention with loop diuretics during the inactive (ZT2) and active (ZT14) phases. Loop diuretics induce similar fractional reductions in oxygen consumption in both sexes: by 9.5% and 8.7% during the inactive phase in the medullary segments of female and male rats, respectively, and by 10.9% and 9.3% during the active phase.

**Figure 8.**
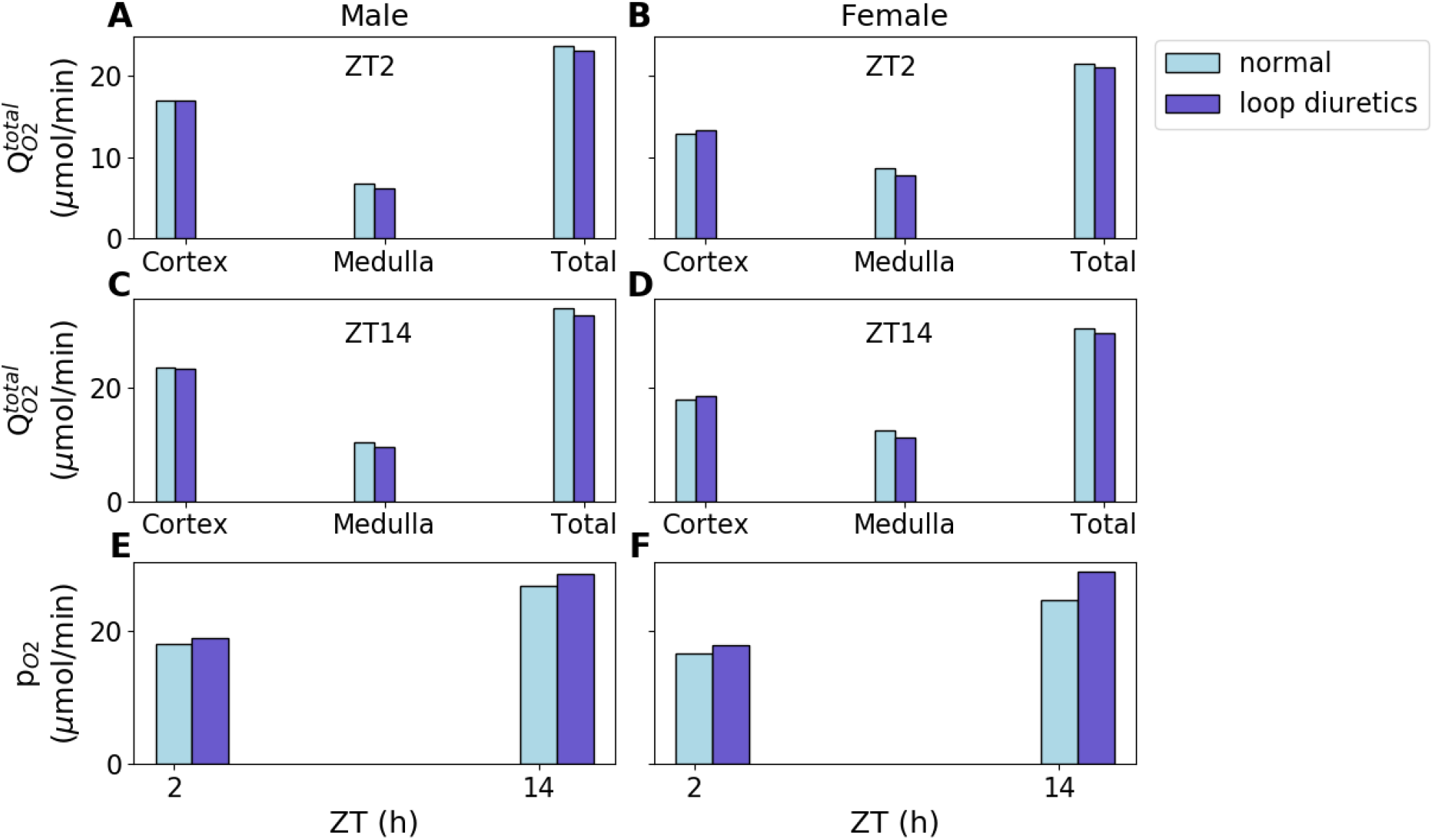
Predicted total before and after inhibition with loop diuretics in the cortical segments, medullary segments, and whole kidney of male and female rats at zeitgeber times 2 h (inactive phase) (A, B) and 14 h (active phase) (C, D). (E, F) Predicted renal medullary oxygen tension before and after inhibition by loop diuretics in male and female rats at zeitgeber times 2 h (inactive phase) and 14 h (active phase).

A reduction in is followed by an increase in, which exhibits much more significant sex difference. For male rats, increased by 4.9% and 6.8% during the inactive and active phases, respectively, whereas, for female rats, the corresponding increases were 6.9% and 17.2%, respectively. Loop diuretics inhibit NKCC2 activity, which in turn lowers Na^+^-K^+^-ATPase activity. Since female rats have higher NKCC2 and Na^+^-K^+^-ATPase activity (two times higher) than male rats, they have higher increase in medullary oxygen tension with loop diuretics. Thus, loop diuretics have significantly greater effect on oxygenation in female rats than in male rats.

## 4. Discussion

Kidney oxygenation is dependent on oxygen delivery and consumption, which in turn are dependent on renal hemodynamics and metabolism, respectively. About 80% of total renal oxygen consumption is attributed to active tubular electrolyte transport [56]. Therefore, renal tissue oxygenation fluctuates with alterations in Na^+^ reabsorption. The kidneys are well perfused but have low oxygen extraction. Thus, the kidneys perform high ATP-requiring transport activities in a low oxygen environment, particularly in the medulla where oxygen perfusion is relatively lower than the cortex. The characteristic vascular architecture and high energy demand to drive tubular solute transport makes the renal medulla especially prone to hypoxia. Oxygen consumption depends on GFR and transporter activities. Since these exhibit diurnal variations and sexual dimorphism, it is important to study the diurnal variations and sexual dimorphism in oxygen consumption to understand how the risk of renal hypoxia varies during the day and between the sexes.

In this study, we developed sex- and time-of-day specific computational models of rat kidney function to assess the diurnal variations in cortical and medullary oxygen consumption and oxygenation in male and female rats. The model predicted significant differences in oxygen consumption during the day and between the sexes. Whole kidney total oxygen consumption was ∼43% higher during the active period relative to the inactive period in both male and female rats, whereas that in the cortical and medullary segments were ∼39% and ∼48% higher during the active period relative to the inactive period. Oxygen consumption also varied between the sexes due to the sexual dimorphism of Na^+^ transporter activities. Male rats showed higher oxygen consumption in the cortical segments (∼1/3 higher), whereas female renal oxygen consumption was higher in the medullary segments (∼1/5 higher).

Renal hypoxia is detected in various kidney diseases and is considered to play an important role in the pathophysiology of both acute kidney injury (AKI) and chronic kidney disease (CKD). Renal hypoxia results in AKI and CKD patients because of decreased renal oxygen supply and increased oxygen consumption. Recent studies have also reported interaction between circadian clocks and hypoxia [57-61]. Circadian clocks control diurnal oscillations in tissue oxygenation, which in turn synchronizes clocks through HIF1α [57, 58]. Clock components simultaneously modulate HIF1α levels and activity [59-61]. A recent study demonstrated that several core clock genes were phase-shifted in response to hypoxia in a tissue-specific manner, which caused inter-tissue circadian clock misalignment [62]. The present study utilized a simple balance equation to predict *p*_O2_. Equation 4 assumes that oxygen supply and consumption are homogenous within the cortical and medullary compartments. However, anatomic studies in the medulla of rodent kidneys have revealed a highly structured organization of nephrons and vessels [63-65]. In the inner stripe, oxygen-carrying descending vasa recta are isolated within tightly packed vascular bundles, separated from the metabolically demanding thick ascending limbs. Simulations using computational models that capture these 3D structures have yielded marked gradients in intrabundle and interbundle interstitial fluid oxygen tension [66-68]. Because of their high metabolic demand and because of their separation from the vascular bundles, the medullary thick ascending limbs operate near hypoxia [69]. The effect of the 3D architecture in the outer medulla on renal oxygenation as well as the heightened risk of hypoxia of the medullary TAL are not captured in the present model.

Loop diuretics are frequently used in the treatment of AKI and CKD. By inhibiting thick ascending limb active Na^+^ transport, loop diuretics have been found to ameliorate medullary hypoxia [48]. Our model predicted that loop diuretics were significantly more effective in improving medullary oxygenation in female rats (by ∼17.2% during the active period compared to the ∼6.8% improvement in male rats). Also, the effect was higher when loop diuretic was administered during the active phase. These results highlight the importance of sex and time of day on physiological functions and drug administration [70]. These factors should be taken into account during biomedical research and modelling analysis. Circadian rhythms can alter the absorption, distribution, metabolism, and excretion rates of drugs. Thus, a fixed dose of drug may result in varying responses depending on the time of administration. Sex and time of day specific computational models can play a major role in improving modern personalized medicine as they can serve as useful tools in determining optimal drug dosages for each sex as well as the most effective time of drug administration.

## Acknowledgements

This study is supported in part by grants from the Natural Sciences and Engineering Research Council (NSERC) and Canadian Institutes of Health Research (CIHR) of Canada to A.T. Layton.

